# The Transition from Acute to Chronic Pain: Dynamic Epigenetic Reprogramming of the Mouse Prefrontal Cortex up to One Year Following Nerve Injury

**DOI:** 10.1101/2020.02.22.956128

**Authors:** Lucas Topham, Stephanie Gregoire, HyungMo Kang, Mali Salmon-Divon, Elad Lax, Magali Millecamps, Moshe Szyf, Laura Stone

## Abstract

Chronic pain is associated with persistent structural and functional changes throughout the neuroaxis, including in the prefrontal cortex (PFC). The PFC is important in the integration of sensory, cognitive and emotional information and in conditioned pain modulation. We previously reported wide-spread epigenetic reprogramming in the PFC many months following nerve injury in rodents. Epigenetic modifications, including DNA methylation, can drive changes in gene expression without modifying DNA sequences. To date, little is known about epigenetic dysregulation at the onset of acute pain or how it progresses as pain transitions from acute to chronic. We hypothesize that acute pain following injury results in rapid and persistent epigenetic remodelling in the PFC that evolves as pain becomes chronic. We further propose that understanding epigenetic remodelling will provide insights into the mechanisms driving pain-related changes in the brain. Epigenome-wide analysis was performed in the mouse PFC 1 day, 2 weeks, 6 months, and 1 year following peripheral injury using the spared nerve injury (SNI) in mice. SNI resulted in rapid and persistent changes in DNA methylation, with robust differential methylation observed between SNI and sham-operated control mice at all time points. Hundreds of differentially methylated genes were identified, including many with known function in pain. Pathway analysis revealed enrichment in genes related to stimulus response at early time points, immune function at later time points and actin and cytoskeletal regulation throughout the time course. Increased attention to pain chronicity as a factor is recommended for both pain research and management.

## 1. Introduction

Chronic pain is a global challenge, affecting between 10-20% of adults and costing billions each year in patient care and lost productivity [21; 29; 53]. Chronic pain is modulated by a complex mix of biological, psychological and social factors. At the molecular level, hundreds of genes become differentially expressed in chronic pain conditions [1; 30; 38; 43; 70]. Understanding chronic pain is further complicated by its development over time from an acute to chronic pain state with time-dependent recruitment of mechanisms and pathways mediating this transition [12; 49]. Improved understanding of pathological gene dysregulation from acute to chronic pain is desperately needed to prevent the development of prolonged, intractable pain.

Epigenetic mechanisms allow for malleable and reversible gene regulation without modification of DNA sequences. DNA methylation is an epigenetic regulator of gene expression wherein a methyl group is added to cytosine DNA nucleotides. When methylation occurs in gene promoter regions, the gene is typically repressed via hindrance of transcription factor binding, recruitment of methyl binding proteins, and chromatin remodeling, ultimately preventing RNA polymerase-mediated transcription [19; 33].

DNA methylation is a stable yet malleable mechanism to regulate gene expression. It is environmentally sensitive, with methylation profiles changing in response to addiction, early life stress, and neurodegenerative disease [46; 51; 54]. Chronic pain is also associated with changes in DNA methylation in both humans and animal models across many tissues including the prefrontal cortex (PFC), dorsal root ganglia, and spinal cord [20; 22; 43; 62]. DNA promoter methylation at specific genes can be linked to the degree of mechanical hypersensitivity [43], and chronic treatment with methyl donor S-adenosylmethionine increased global DNA methylation in the PFC and attenuated mechanical hypersensitivity [23]. Investigation of DNA methylation’s role in chronic pain advances our understanding of pain progression and may reveal potential therapeutic avenues.

Among the structures involved in chronic pain, the prefrontal cortex (PFC) is crucial as an integrator of ascending sensory information, cognitive and emotional responses, and descending inhibitory control [10; 37; 69]. In human patients and animal subjects, long-term pain results in a persistent functional and anatomical reorganization of the PFC [5; 36; 56; 58], with patients showing more PFC activation than healthy controls [4; 6]. Chronic pain patients are commonly comorbid for depressive and anxiety disorders [57] that are heavily correlated with PFC abnormal functioning [34; 50]. As a central mediator in the pain pathway, therapeutic mechanisms targeting PFC may have efficacy across different pain disorders and simultaneously address associated cognitive and emotional comorbidities.

Previous work has shown extensive changes in DNA methylation and dynamic structural and functional reorganization in the PFC as a result of chronic pain. To track epigenetic reprogramming in the PFC in acute and chronic pain, we captured the pattern of genome-wide DNA methylation across time from 1 day to 1 year following peripheral nerve injury in mice. Our analysis revealed a dynamic pattern of DNA methylation across the time course, implicating hundreds of differentially methylated genes at each time point. Subsequent analysis revealed time point-specific changes that reveal potential underlying mechanisms.

## 2. Materials and Methods

### 2.1 Animals

78 male CD-1 mice (Charles River Laboratories, St-Constant, QC, Canada) mice were used in this study. Animals were received at 6-8 weeks of age and housed 3-4 per cage on a 12h light/dark cycle in a temperature controlled room in ventilated polycarbonate cages (Allentown, Allentown, NJ) with corncob bedding (7097, Teklad Corncob Bedding, Envigo, UK) and cotton nesting squares for enrichment. Mice were given access to food (2092X Global Soy Protein-Free Extruded Rodent Diet, Irradiated) and water *ad libitum*. Animals were habituated to the housing conditions for at least one week prior to any experimental interventions.

Animals designated for DNA methylation analysis were randomly assigned to receive either the spared nerve injury (SNI) model of neuropathic pain or sham surgery control and the model was allowed to develop for one day (D1), two weeks (W2), six months (M6), or one year (1Yr) post-injury. Final numbers per group (Sham/SNI) – D1:4/4; W2:6/6; M6:6/6; 1Yr:4/6; 42 animals in total.

Animals designated for immunohistochemistry were randomly assigned to either SNI or sham surgery and allowed to progress to two weeks or six months post-injury. Final numbers per group (Sham/SNI) – W2:10/8; M6:10/8; 36 animals in total.

All experiments were approved by the Animal Care Committee at McGill University, and conformed to the ethical guidelines of the Canadian Council on Animal Care and the guidelines of the Committee for Research and Ethical Issues of the International Association for the Study of Pain [75].

### 2.2 Induction of spared nerve injury

Nerve injury was induced using the Spared Nerve Injury model of neuropathic pain, as adapted for mice [14; 59] at 10 to 12 weeks of age. Under deep isoflurane anesthesia, an incision was made on the medial surface of the thigh, exposing the three branches of the sciatic nerve. SNI surgery consisted of ligation and transection of the left tibial and common peroneal branches of the sciatic nerve, while sparing the sural nerve. The tibial and common peroneal branches were tightly ligated with 6:0 silk (Ethicon) and sectioned distal to the ligation. Sham surgery consisted of exposing the nerve without damaging it.

### 2.3 Behavioural assessment of mechanical sensitivity

After a habituation period of 1 hour to the testing environment of Plexiglas boxes placed on a wire mesh grid, von Frey filaments (Stoelting Co, Wood Dale, IL) were applied to the plantar surface of the hind paw to the point of bending for 3 seconds or until withdrawal. Mechanical sensitivity was determined as the 50% withdrawal threshold using the up-down method [11]. The stimulus intensity ranged from 0.04 to 4.0 g, corresponding to filament numbers 2.44 to 4.56. Testers were blind to the condition of the animal.

### 2.4 Immunofluorescent histochemistry

Animals were deeply anesthetized with an intraperitoneal injection of ketamine (100 mg/kg), xylazine (10 mg/kg), and acepromazine (3 mg/kg). Animals were perfused with 100 mL vascular rinse (0.2M phosphate buffer with sodium nitrate), followed by 200 mL Lana’s fixative (4% paraformaldehyde, 14% saturated picric acid, 0.16M phosphate buffer, pH 6.9), and then 100 mL 10% sucrose solution in phosphate buffered saline. Tissue was cryoprotected in 10% sucrose solution at 4°C for three days, and then coronally divided at Bregma into forebrain and hindbrain sections. Brain tissue was coronally sectioned at 14μm on a cryostat (Leica), and PFC sections, defined as +1.54 mm to +1.98 mm from Bregma, were collected. Prelimbic and infralimbic areas were defined according to Paxinos and Franklin [52].

Tissue was incubated with blocking buffer (1% Normal Donkey Serum, 1% Bovine Serum Albumin, 0.3% Triton X-100, 0.01% Sodium Azide) for 1 hour before primary antibody incubation overnight at 4°C. Tissue sections were either singly stained with rabbit anti-NeuN (1:1000, Abcam, Ab177487), or double stained with mouse anti-NeuN (1:1000, Millipore, MAB377) and either rabbit anti-GFAP (1:1000, Neomarkers, RB087AO) or rabbit anti-Iba1 (1:1000, Wako, 019-19741). Rabbit anti-NeuN was used to quantify neuronal cell counts while mouse anti-NeuN was used as a counter stain to define cortical layers in double-stained sections. Secondary antibodies (donkey anti-rabbit Alexa 594 (1:200, Jackson ImmunoResearch, 711-585-152) and donkey anti-mouse Alexa 488 (Jackson ImmunoResearch, 714-545-150) were incubated for 90 min at room temperature. Sections were counterstained with DAPI (1:50000, Sigma, D9542) and coverslipped with AquaPolymount (18606, PolySciences).

### 2.5 Immunohistochemistry quantification

FIJI [55] and manual cell counting were used to quantify the number of total cells, neurons, astrocytes, and microglia in prelimbic and infralimbic brain areas. DAPI was used as a marker of total cell population, NeuN-immunoreactivity (-ir) defined neuronal populations, GFAP-ir defined astrocyte populations, and Iba1-ir defined microglial populations. NeuN staining was also used to separately designate layer I, superficial (layers II-III) and deep (layer V) cortical layers to aid the placement of three regions-of-interest (ROIs) per layer. Cell quantification was based on colocalization of the cell type specific marker with DAPI, and cell counts were averaged across three ROIs per layer. The averaged ROI area per cortical layer: layer I: 6724 µm^2^; superficial (layers II-III): 15790.14 µm^2^, deep (layer V): 10526.76 µm^2^. An overall cell count was calculated by adding the counts from all three layers. Total ROI area: 33040.9 µm^2^. ROI sizes were identical between prelimbic and infralimbic areas.

### 2.6 Isolation of prefrontal cortex, DNA capture and bisulfite sequencing

Mouse PFC was dissected following isoflurane anesthesia and decapitation. PFC was defined as a region with an anterior boundary of the frontal pole, approximately +3.00 Bregma, posterior boundary of +1.42 Bregma, lateral boundaries of +1 mm from midline, and a ventral boundary wherein the olfactory bulb, caudate putamen, and areas below the rhinal fissure and lateral ventricles were all removed [52]. This defined area isolates the prelimbic, orbital, and infralimbic cortices corresponding to the mouse medial PFC. Once dissected, tissue was homogenized and genomic DNA was extracted using the Qiagen DNeasy Blood and Tissue kit and protocol.

Genome-wide methylation analysis was performed using the Roche SeqCap Epi Developer M Enrichment system on bisulfite-treated DNA (Roche Nimblegen; WI, USA), with custom designed primers targeting promoter and enhancer regions of the genome. Probe design was based on the mm9 reference genome and associated H3K4me1 and H3K4me3 binding (probe BED file available in **Supplemental Table 1**). Briefly, 1 µg of genomic DNA per sample was fragmented into 200bp lengths, and adapter and index sequences were ligated to each end of the fragment according to the KAPA Library Preparation Kit protocol. Sample DNA was then bisulfite converted following the EZ DNA Methylation-Lightning Kit (Zymo Research; CA, USA). Sample DNA was then PCR amplified, hybridized with custom capture probes, and PCR amplified again in accordance with Roche SeqCap Epi system protocols [68]. Sample captured bisulfite-converted DNA sequencing was performed by Genome Quebec on the Illumina HiSeq 2500 following Illumina guidelines.

### 2.7 DNA methylation pre-processing

Following Illumina Sequencing, sample data was received as FASTQ files and organized as 125bp paired end reads. Capture sequencing data was pre-processed using the McGill University Genome Quebec Innovation Centre GenPipes Methyl-Seq Pipeline [8]. The pipeline proceeds through Trimmomatic, Bismark Align, Picard Deduplication, and Bismark Methylation Call [7; 35]. Baseline pipeline parameters were set as a minimum sequence quality of PHRED score > 30, alignment reference genome was mm10 (GRCm38), and minimum read coverage per base was set at 10 reads. A detailed outline of the Methyl-Seq pipeline steps and process can be found at https://bitbucket.org/mugqic/genpipes. Before analysis, collected sequence data was filtered to remove a blacklist of regions identified to have anomalous, unstructured, and high signal/read counts in next gen sequencing [2]. The blacklist used was specific for mm10 and the BED file can be found in **Supplemental Table 2**.

### 2.8 Differential methylation analysis

Generalized linear model analysis was used to compare Sham to SNI methylation values at each time point. Differential methylation analysis was performed using R and the edgeR package [13]. Samples were normalized to library size, where library size was set to be the average of the total read counts for methylated and unmethylated CpGs. For a CpG site to pass through to analysis, it was required to be sequenced at a depth of at least 10 reads in each sample within a time point.

At each time point, CpGs were defined as being differentially methylated if significance was calculated to be FDR adjusted p-value < 0.1. While the selection of an FDR threshold of 0.1 carries greater risk of type I error than the more typical threshold of 0.05, it is preferable for the exploratory purposes of this study. Differential methylation between Sham and SNI was determined by calculating the M-Value (log_2_ of the ratio (methylated cytosines / unmethylated cytosines)) of each CpG site in Sham and SNI, and then detecting statistically significant differences. The methylation percentage (percentage of (methylated cytosines / total cytosines)) at each site was also calculated. Differentially methylated samples were further divided into hypermethylated CpGs or hypomethylated CpGs with the Sham CpG methylation percentage acting as the reference point. If the SNI CpG methylation is 5% or larger than the Sham, the CpG position is considered to be hypermethylated. If the SNI CpG methylation is reduced by −5% or more compared to Sham, the CpG position is considered to be hypomethylated.

For gene annotation, CpGs were annotated to their associated genes and genomic regions using the topTags function of edgeR [13] and annotatePeaks function of the ChIPseeker [72] R packages, using a TxDb object created from the Gencode “mmusculus_gene_ensembl” Biomart database. Each CpG position was annotated and tracked individually throughout the analysis. For homology conversion, gene symbols from non-mouse species were converted to known mouse homologs based on the Mouse Genome Database (MGD) – Vertebrate Homology, Mouse Genome Informatics, The Jackson Laboratory, Bar Harbor, Maine [9].

### 2.9 Statistical analysis

Two-tailed Student’s t-test tested for group differences between Sham and SNI in the von Frey mechanical sensitivity behavioural assay (Figure 1). A two-tailed Welch’s t-test was used to test for differences in cell type counts between Sham and SNI (Figure 2). Differences between the magnitude of differential methylation across time points was identified using a one-way ANOVA with a Bonferroni correction for multiple comparisons (Figure 3). Pairwise comparisons of CpG positions at each time point utilized a negative binomial generalized linear model to fit the data and differential methylation was determined utilizing the likelihood ratio test. Analyses were corrected for multiple comparisons using the Benjamini-Hochberg false discovery rate (Figure 4). Post-hoc supervised hierarchical clustering analysis was performed using Ward’s method and 1 – Pearson correlation as the distance metric (Figure 5-8A).

**Figure 1.**
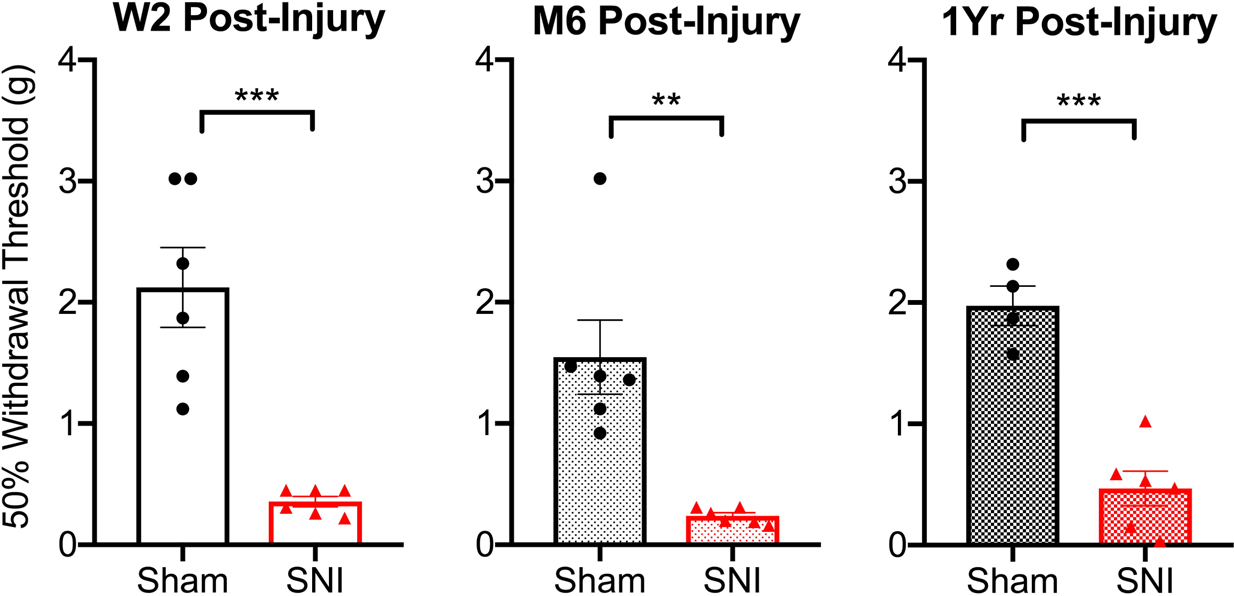
Mechanical hypersensitivity persists for up to 1 year following spared nerve injury (SNI). Nerve injured mice were hypersensitive to mechanical stimuli compared to sham mice at two weeks (W2), six months (M6), and one year (1Yr) post-SNI. Bars represent mean ± SEM withdrawal threshold expressed in grams. *Two-tailed Student’s t-test,**=p<0.01; ***=p<0.001, n=4-6/group*.

**Figure 2.**
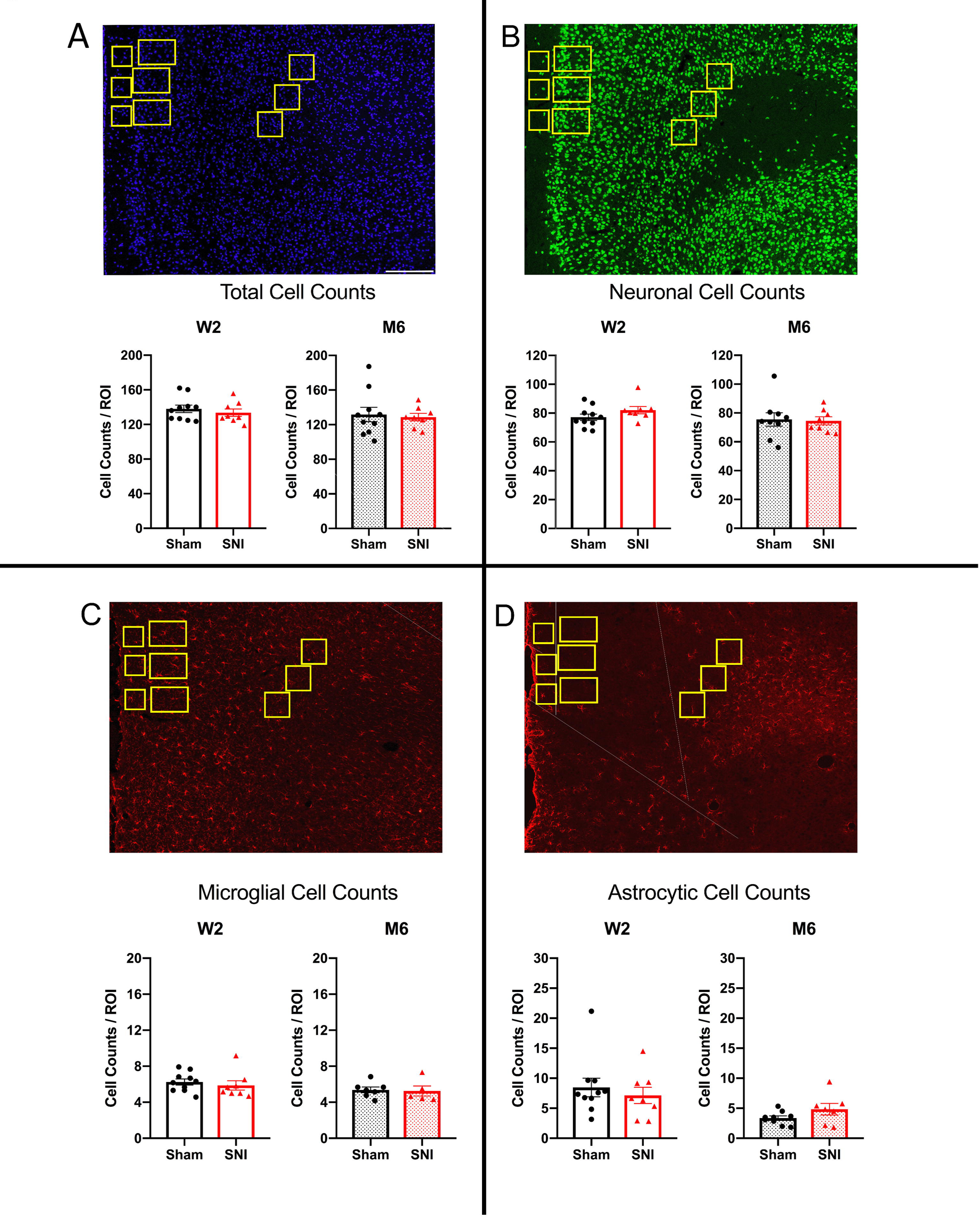
Immunohistochemistry reveals no changes in neuronal, microglial or astrocytic cell proportions two-weeks and six-months following spared nerve injury (SNI). A) Total (DAPI, Blue), B) Neuronal (NeuN-immunoreactivity (-ir), Green), C) microglial (Iba1-ir, Red), and D) astrocytic (GFAP-ir, Red) cell counts in Sham and SNI cortex at W2 or M6 time points. Displayed are average cell counts from all cortical layers from prelimbic area of the prefrontal cortex. Quantified ROI area is equivalent to 33040.9 µm^2^. Bars represent mean ± SEM cell counts; *Two-tailed Welch’s t-test. No significant differences observed. n = 8-10/group*.

**Figure 3.**
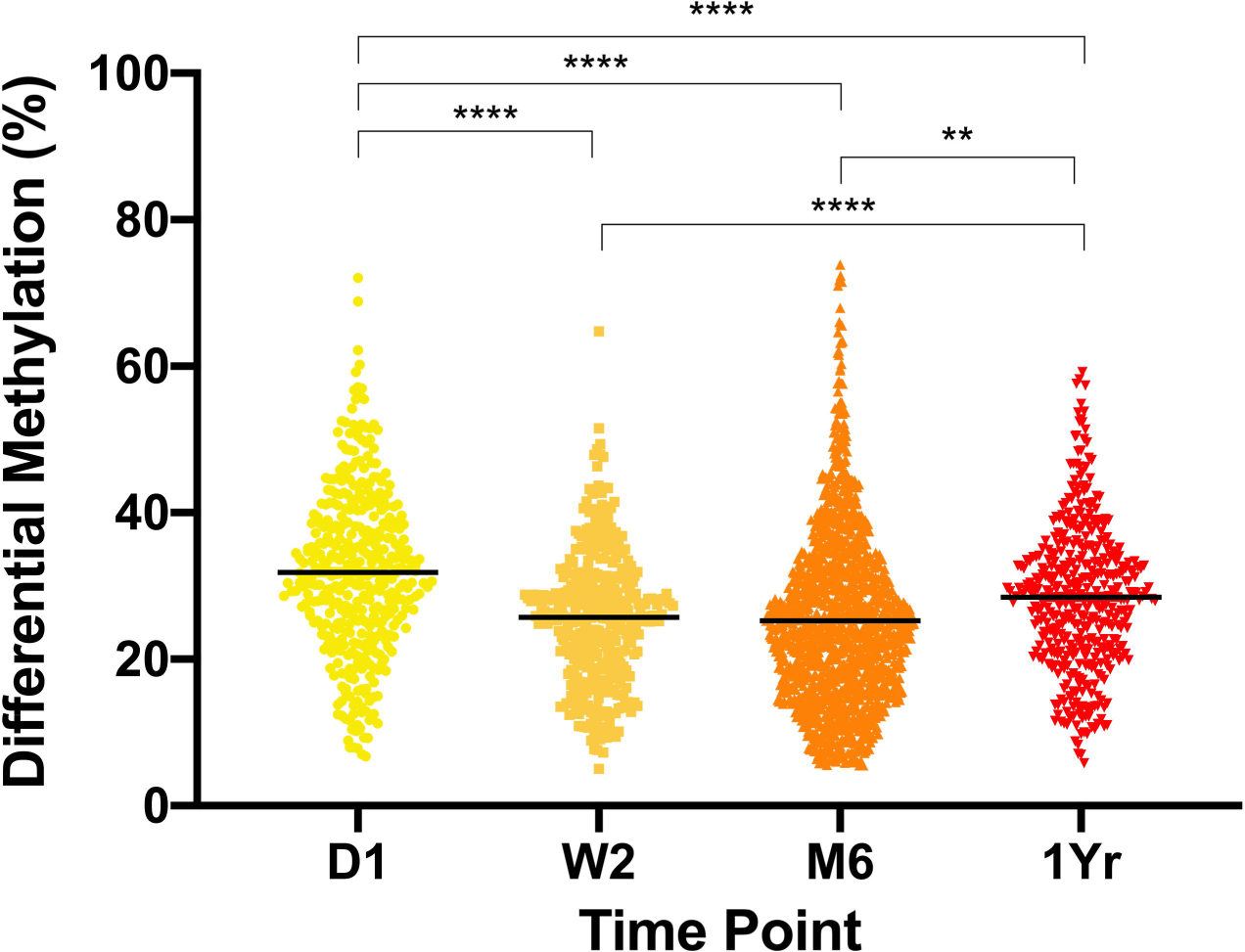
Magnitude of differential methylation following spared nerve injury (SNI) Differential methylation is expressed in terms of absolute percentage of change between Sham and SNI at 1 day (D1), 2 weeks (W2), 6 months (M6) and 1 year (1Yr) post-surgery. Each dot is an individual CpG. *Welch’s ANOVA with Games-Howell’s test for multiple comparisons.*. *\*\*p<0.01, ****p< 0.0001. n = 4-8/group*.

**Figure 4.**
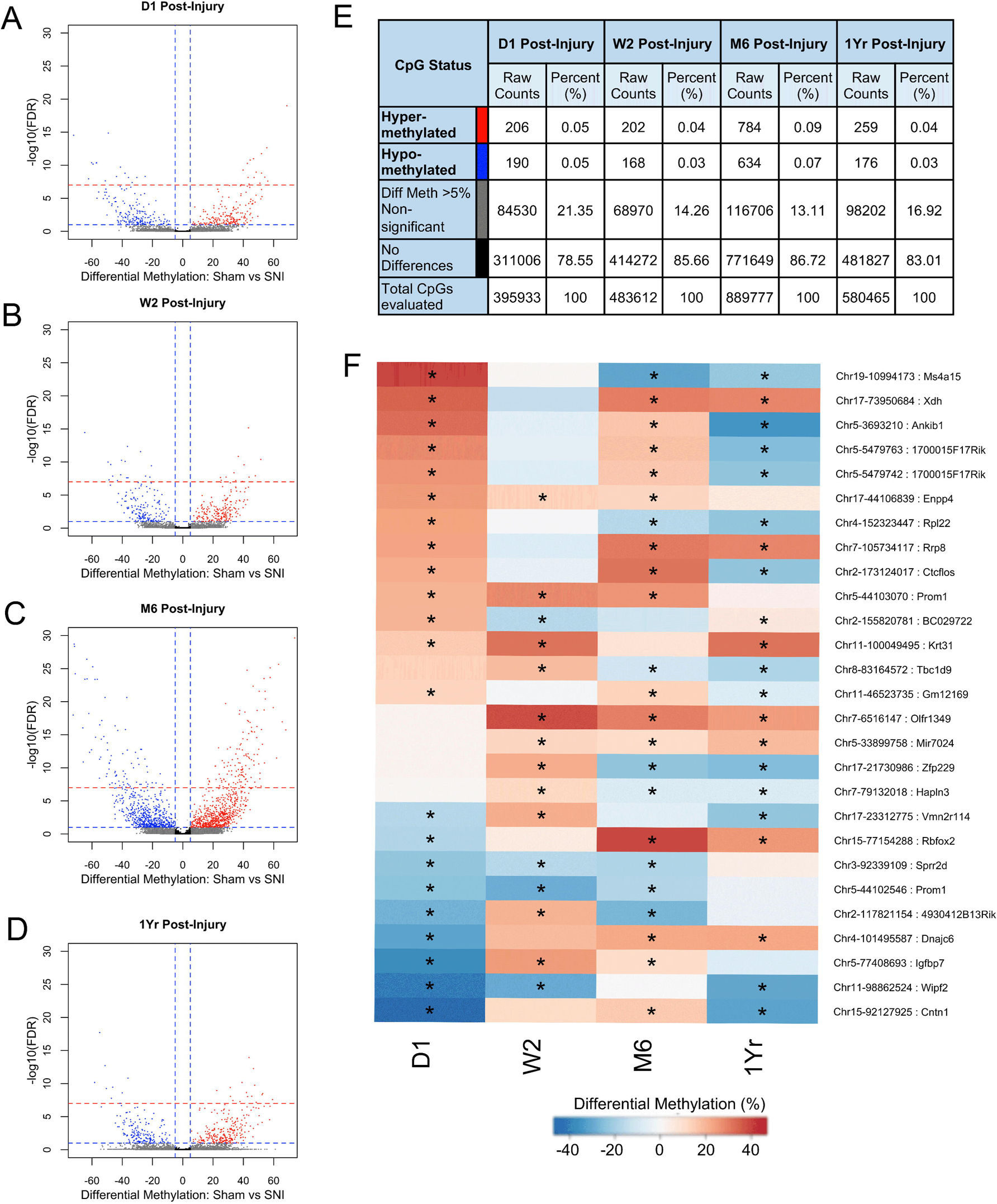
Hyper- and hypo-methylated CpGs following spared nerve injury (SNI). The magnitude of the statistical significance between methylation differences in promoter region CpGs at A) 1 day (D1), B) 2 weeks (W2), C) 6 months (M6) and D) 1 year (1Yr) post-surgery. CpGs are displayed in terms of positive or negative methylation difference with Sham as the reference point vs. SNI, against –log10(FDR) on the y-axis. Blue horizontal dashed line indicates an FDR adjusted p-value threshold of 0.1, red horizontal dashed line indicates an FDR adjusted p-value threshold of 1×10^-7^. The blue vertical dashed line indicates methylation difference thresholds of 5% and −5%. Red dots = hyper-methylated CpGs: blue dots = hypo-methylated CpGs; grey dots = non-significant with methylation differences of >5%, black dots = CpGs that are not different: black. E) The raw counts and percentage of detected CpGs. F) Single CpG positions that were significantly different at three or more time points were tracked across time and display varying patterns of differential methylation. Values are displayed as differential methylation. *\* = FDR adjusted p-value < 0.1*.

**Figure 5.**
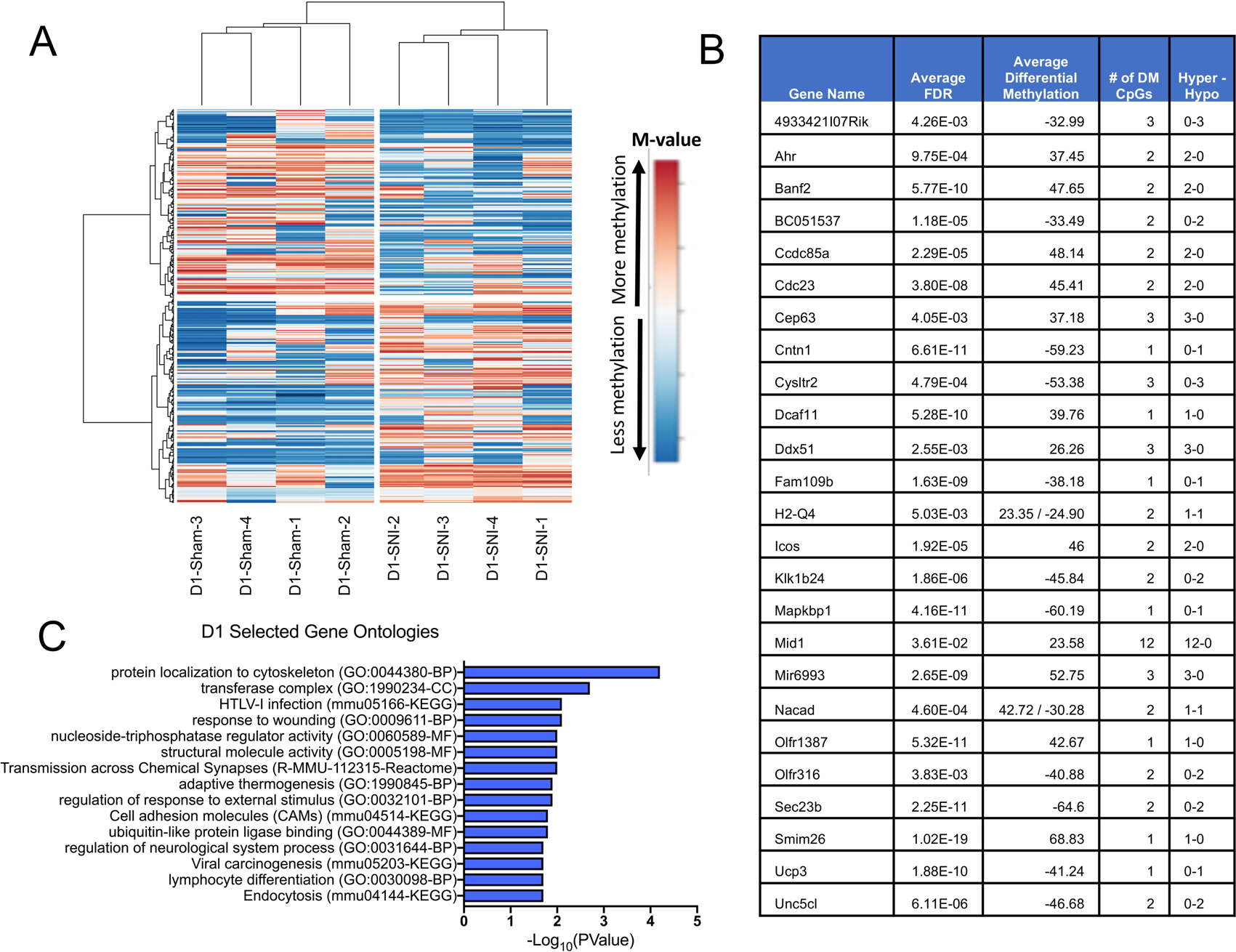
Differentially methylated genes and gene ontologies at 1 day (D1) following spared nerve injury (SNI). **A)** Post-hoc supervised hierarchical clustering of DM CpGs at each time point displays common patterns of differential methylation between individual animals in the same treatment group. Values are displayed as M-Value: log_2_(# methylated CpGs / # unmethylated CpGs) Clustering method: Ward, distance metric: 1-Pearson correlation. **B)** Top 25 differentially methylated genes, **C)** Top 15 selected gene ontologies identified. *Enriched ontology: minimum gene count of 3, p < 0.05, and a minimum enrichment of 1.5. n = 4 mice/group*.

### 2.10 Gene ontology analysis

Differentially methylated genes identified at each time point were submitted to Metascape [74], a web-based Gene Ontology tool. The CORUM, the Reactome Gene Sets, the KEGG Pathway, Functional Set and Structural Complex, and the GO Biological Processes, Cellular Component and Molecular Function databases were selected as ontology databases. An ontology was considered enriched at a p-value < 0.05, minimum gene count of 3, and a minimum enrichment factor > 1.5.

## 3. Results

### 3.1 Spared nerve injury results in long-term neuropathic pain but has no impact on the proportions of neurons, astrocytes and microglia in the prefrontal cortex

Before performing bisulfite sequencing, we validated the presence of a long-term neuropathic pain (Figure 1). To control for non-epigenetic mechanisms that might drive differential DNA methylation, we verified that the proportions of neurons, astrocytes and microglia do not differ between Sham and SNI animals at W2 and M6. As each cell type has its own DNA methylation profile, differential methylation between Sham and SNI animals could result from the loss of specific cell populations.

#### 3.1.1 Behavioural signs of neuropathic pain persist up to one year following spared nerve injury

To confirm long-term neuropathic pain, mechanical sensitivity was assessed using the von Frey assay at 2 weeks, 6 months, and 1 year following SNI. SNI animals were more sensitive to mechanical stimuli applied to the plantar surface of the hindpaw ipsilateral to the injury at all time points when compared to their respective sham animals (Two-tailed Student’s t-test. *W2*: t_10_ = 5.3, p = 0.0003; *M6*: t_10_ = 4.2, p = 0.002; *1Yr:* t_8_ = 6.8, p = 0.0001) (**Figure 1**). The reduction in 50% withdrawal thresholds in SNI animals indicates the presence of a persistent and stable mechanical hypersensitivity for up to 1 year.

#### 3.1.2 Spared nerve injury has no impact on the proportions of neurons, astrocytes and microglia in the prefrontal cortex

The neuronal, astrocytic, and microglial cell populations were quantified in the PFC by immunohistochemistry to quantify SNI-related changes 2 weeks and 6 months following surgery. Overall cell counts were calculated independently in prelimbic and infralimbic cortical areas as well as at three cortical depths: layer I, superficial layers (layer II-III), and deep layers (layer V). No significant differences were detected between SNI and Sham for any cell types in either the prelimbic (Two-tailed Welch’s t-test. **DAPI** *W2*: t_15.80_ = 0.7 p = 0.48; *M6*: t_13.21_ = 0.3, p = 0.75. **NeuN** *W2*: t_15.16_ = 1.5, p = 0.16; *M6*: t_12.96_ = 0.2, p = 0.86. **GFAP** *W2*: t_16_ = 0.6, p = 0.53; *M6*: t_7.80_ = 1.4, p = 0.19. **Iba1** *W2:* t_12.48_ = 0.6, p = 0.55; *M6:* t_6.51_ = 0.2, p = 0.86) (**Figure 2**) or infralimbic (Supplemental Figure 1) regions at either time point.

### 3.2 Spared nerve injury produces rapid, large and persistent changes in DNA methylation in the prefrontal cortex

Differentially methylated (DM) CpG sites were identified by bisulfite capture sequencing by comparing SNI vs. Sham. In this study only DM CpGs that were annotated to promoter regions were selected for analysis due to the inverse relationship between CpG DNA methylation and gene expression (in general, less methylation = increased gene expression; more methylation = reduced gene expression [67]) DM CpGs are defined as having an FDR adjusted p-value of < 0.1 and a difference in methylation between Sham and SNI of > 5%.

**Table 1** shows the number of DM CpG sites after adjusting for false discovery rate (FDR). At each of the four time points evaluated, at least 350 DM CpGs were identified at an FDR adjusted p-value threshold of 0.1. Increasingly stringent thresholds still identify a large number of differentially methylated positions. Notably, a subpopulation of DM CpGs remain significant at an FDR adjusted p-value < 1×10^-7^, representing a substantial and consistent change between Sham and SNI and strong effect of injury.

**Table 1.** Spared nerve injury (SNI) results in highly significant differentially methylated (DM) CpG positions in prefrontal cortex. SNI induces DM CpGs at all time points. Bolded figures indicate DM CpGs at FDR adjusted p-value experimental threshold of 0.1. *Likelihood ratio test with Benjamini-Hochberg false discovery rate correction for multiple comparisons*.

| FDR | D1 | W2 | M6 | 1Yr |
| --- | --- | --- | --- | --- |
| 1.0 x 10<sup>-7</sup> | 31 | 28 | 214 | 25 |
| 1.0 x 10<sup>-6</sup> | 43 | 42 | 259 | 41 |
| 1.0 x 10<sup>-5</sup> | 62 | 58 | 344 | 62 |
| 1.0 x 10<sup>-4</sup> | 87 | 91 | 424 | 100 |
| 0.001 | 120 | 141 | 605 | 156 |
| 0.01 | 205 | 232 | 883 | 257 |
| 0.1 | 396 | 370 | 1418 | 435 |
| 0.2 | 508 | 487 | 1767 | 541 |

The magnitude of differential methylation at each time point of all DM CpGs is displayed in **Figure 3**. The average magnitude of differential methylation between Sham and SNI is approximately 25-30% at all time points. This is a robust shift in methylation state for most DM CpG positions and is on par with that observed in the brain for major depressive disorder [32].

The effect of peripheral injury on prefrontal cortex methylation is immediate, with the largest degree of differential methylation between Sham and SNI detected at D1 post-injury, indicating a rapid response to injury. D1 had a significantly higher average methylation change between Sham and SNI than the other time points (Welch’s ANOVA, Games-Howell’s multiple comparisons test: D1 vs. W2/M6/1Yr, p < 0.0001), with 75% of CpGs having a greater than 25% change in methylation, and 25% of CpGs having a change larger than 38%. The W2 and 1Yr time points were the most tightly clustered, with 50% of DM CpGs within 6-7% of their mean differential methylation. Despite having the lowest average percent differential methylation, the M6 time point had a large subpopulation skewed towards high levels of methylation change. These highly DM CpGs may play an important role in the transition from subacute to chronic pain.

The effect of injury upon PFC methylation is also persistent, with differential methylation observed throughout the time course (**Figure 4A-D**, dashed blue line: FDR adjusted p-value = 0.1, dashed red line: FDR adjusted p-value = 1×10^-7^). Within each time point there were no significant differences in the number of magnitude of hyper-vs. hypo-methylation (**Figure 4A-D**, red and blue points). This balance suggests locus-specific methylation targeting as opposed to a global increase or decrease in DNA methylation (**Figure 4E**).

To explore how patterns of methylation evolve at individual CpGs across time, we grouped all CpGs that were differentially methylated at a minimum of 3 out of the 4 time points (**Figure 4F**). While for some CpGs the direction of differential methylation is largely consistent across time, others display dynamic patterns as a function of chronicity. For example, the actin cytoskeleton remodeling gene, WAS/WASL interacting protein family member 2 (*Wipf2*), and the cell adhesion gene Contactin 1 (*Cntn1*), are both differentially methylated across the entire time course. Taken together, the DNA methylation response to peripheral nerve injury is rapid, large, dynamic, and persists throughout the entire time-course.

### 3.3 Spared nerve injury results in time point-specific differential methylation of individual genes and functional pathways in the prefrontal cortex

To determine the functional impact of these substantial shifts in the PFC methylation landscape, we performed gene annotation and post-hoc analyses to identify associated genes and gene ontologies.

DM CpGs were annotated to their corresponding gene based on the mm10 reference genome and the Gencode “mmusculus_gene_ensembl” Biomart database. 61.3% of annotated CpGs (1604 / 2619 total DM CpGs) were the only DM CpG annotated to a gene promoter within a time point. In contrast, 35.5% of annotated CpGs (931 / 2619) were part of clusters of 2-6 DM CpGs annotated to a promoter region. The remaining 3.3% of CpGs (84 / 2619) existed within clusters of 7+ CpGs, located to the promoters of 4 genes*, Organic Solute Carrier Partner 1 (Oscp1), Midline1 (Mid1), Toll-like receptor 6 (Tlr6) and 1110020A21Rik*.

Of the genes identified with multiple DM CpGs (349 / 394) within a time point, the vast majority (88.5%) were either all hyper- or all hypo-methylated within that promoter, suggesting that the resulting dysregulated genes are likely to be either up- or down-regulated. In contrast, the remaining genes (45 / 394) had mixed differential methylation in their promoter region, reflecting the complex relationship between promoter methylation and specific regulation of transcription factors with methylation-sensitive binding.

Post-hoc supervised clustering was performed at each time point to visualize clusters of covarying DM CpGs and the characteristic methylation profile of Sham and SNI animals (**Figure 5A-8A**). Within each time point, each row represents a CpG position, with samples coloured according to the M-Value measure of methylation, with blue being less methylation and red being more methylation.

To identify the most differentially methylated genes at each time point, a promoter region index was determined by multiplying the adjusted p-value of each CpG by the number of CpGs within the promoter region. The top 25 differentially methylated genes at each time point as determined by promoter region index can be viewed in **Figures 5B-8B**, listing the average adjusted p-value, average differential methylation, number of DM CpGs identified within the gene promoter, and number of hyper- or hypo-methylated DM CpGs. In cases with mixed differential methylation in the promoter, the average methylation for both hyper-methylated and hypo-methylated CpGs are shown separately. Full lists displaying individual DM CpGs and their annotated genes are available in **Supplemental Table 3**.

To identify broader effects of differential gene methylation and therefore potential functional dysregulation, gene ontologies enriched for differentially methylated genes were also identified (**Figure 5C-8C**). At each time point, 15 representative gene ontologies were selected on the basis of p-value, number of differentially methylated genes within the ontology, and being a parent term to other identified ontologies. At most, only 66% of differentially methylated genes within an ontology are identical to the differentially methylated genes identified in any other ontology at each time point. The full list of enriched ontologies by time point can be observed in **Supplemental Table 4**.

### 3.4.1 One day post-injury

At D1, 396 DM CpGs identified 327 unique genes undergoing differential methylation in their promoter region. The top 25 differentially methylated genes were related to cell-cell interaction (*Ccdc85a, Cntn1*), cell structure (*Banf2, Cep63, Mid1*), and immune response (*Cystlr2, H2-Q4, Icos, Mapkbp1, Unc5cl*) (**Figure 5B**). *Mapkbp1, H2-Q4, Icos*, *Unc5cl*, have involvement in TLR, NFκB, and TNF signaling pathways, with additional immune related genes *Acod1, Cd14, Itch, H2-K1, H2-M1, Mapkapk3, Mapkapk5,* and *Tnfrsf13c* also identified as differentially methylated at this time point. While not part of the top 25 differentially methylated genes, actin and cytoskeletal dynamics genes (*Acta2, Diaph1, Dvl1, Macf1, Wipf2*) are also observed.

Gene ontology analysis identified functional pathways enriched for differentially methylated genes involved in response to stimulus, cellular structure remodeling, and extracellular matrix formation. Specifically, top gene ontologies included: GO:0044380 – Protein Localization to Cytoskeleton (p = 6.31×10^-5^, 6 differentially methylated genes), transferase complex – (p = 1.99×10^-3^, 19 genes), GO:0009611 – Response to Wounding (p = 7.94×10^-3^, 13 genes), GO:0005198 – Structural Molecule Activity (p = 0.01, 15 genes), GO:0032101 – Regulation of Response to External Stimulus (p = 0.013, 17 genes), and mmu04514 – Cell Adhesion Molecules (p = 0.012, 6 genes) (**Figure 5C**).

### 3.4.2 Two weeks post-injury

At W2, 370 DM CpGs identified 311 unique genes undergoing differential methylation in their promoter region. The top 25 differentially methylated genes had involvement in actin processes (*Mical3*), cell surface molecules (*Gpc6, Prom1*), and immune regulation (*Btla, Ccl7*) (**Figure 6B**), In addition to the top 25, the W2 time point revealed a relatively large number of genes involved in cellular structure and organization processes, such as actin and cytoskeleton (*Capza3, Epb41l1, Fhdc1, Myh2, Mprip, Shroom3, Tns1, Tns4, Wipf2)*, extracellular matrix (*Col4a1, Col6a4, Halpn3, Rpsa*) and cell-cell interactions (*Nectin2, Lgals4*). Also of interest is the differential methylation of *Ramp1,* which is crucial for the maturation and transport of CALCRL and its function as the CGRP receptor.

**Figure 6.**
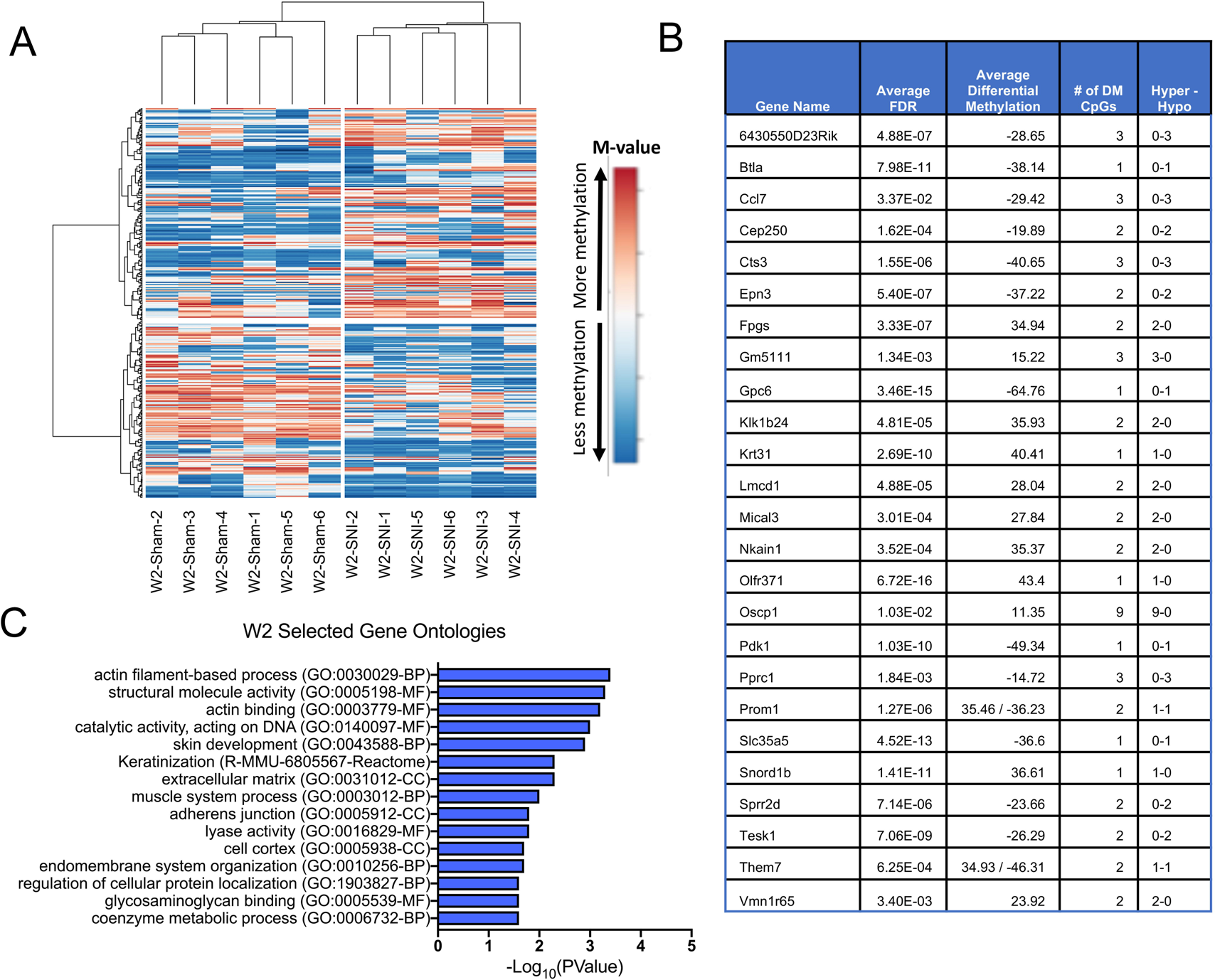
Differentially methylated genes and gene ontologies 2 weeks (W2) following spared nerve injury (SNI). **A)** Post-hoc supervised hierarchical clustering of DM CpGs at each time point displays common patterns of differential methylation between individual animals in the same treatment group. Values are displayed as M-Value: log_2_(# methylated CpGs / # unmethylated CpGs) Clustering method: Ward, distance metric: 1-Pearson correlation. **B)** Top 25 differentially methylated genes, **C)** Top 15 selected gene ontologies identified. *Enriched ontology: minimum gene count of 3, p < 0.05, and a minimum enrichment of 1.5. n = 6 mice/group*

At W2, identified gene ontologies are involved with cellular structure remodeling, extracellular matrix, response to stimulus, and changes to internal cellular functioning. Specifically, top ontologies included: GO:0030029 – Actin Filament-Based Process (p = 3.98×10^-4^, 19 genes), GO:0005198 – Structural Molecule Activity (p = 5.01×10^-4^, 17 genes), GO:0003779 – Actin Binding (p = 6.31×10^-4^, 13 genes), GO:0031012 – Extracellular Matrix (p = 5.01×10^-3^, 12 genes), GO:1903827 – Regulation of Cellular Protein Localization (p = 0.019, 11 genes)(**Figure 6C**).

### 3.4.3 Six months post-injury

At M6, 1418 DM CpGs identified 1001 unique genes undergoing differential methylation in their promoter region. The top 25 differentially methylated genes were involved in E3 ubiquitination (*Mid1, Trim21, Trim30d*), and immune functioning (*Gdf15, Fam129c, Arhgef2*) (**Figure 7B**), In addition to the top 25, the M6 time point also displays a substantial number of immune genes across many different pathways (*Cd6, Ltbp1, Trem2;* TNF: *Traf3ip1, Traf6;* Interleukin: *Il12b, Il12rb1, Jak3;* TLR: *Lbp, Acod1, Myd88, Tlr6*). Actin regulation and formation is also affected, with numerous differentially methylated GTPases and guanine exchange factors (*Arhgap8, Arhgap15, Arhgap17, Arhgap33os, Arhgef2, Arhgef10l, Arhgef17, Arhgef19*).

**Figure 7.**
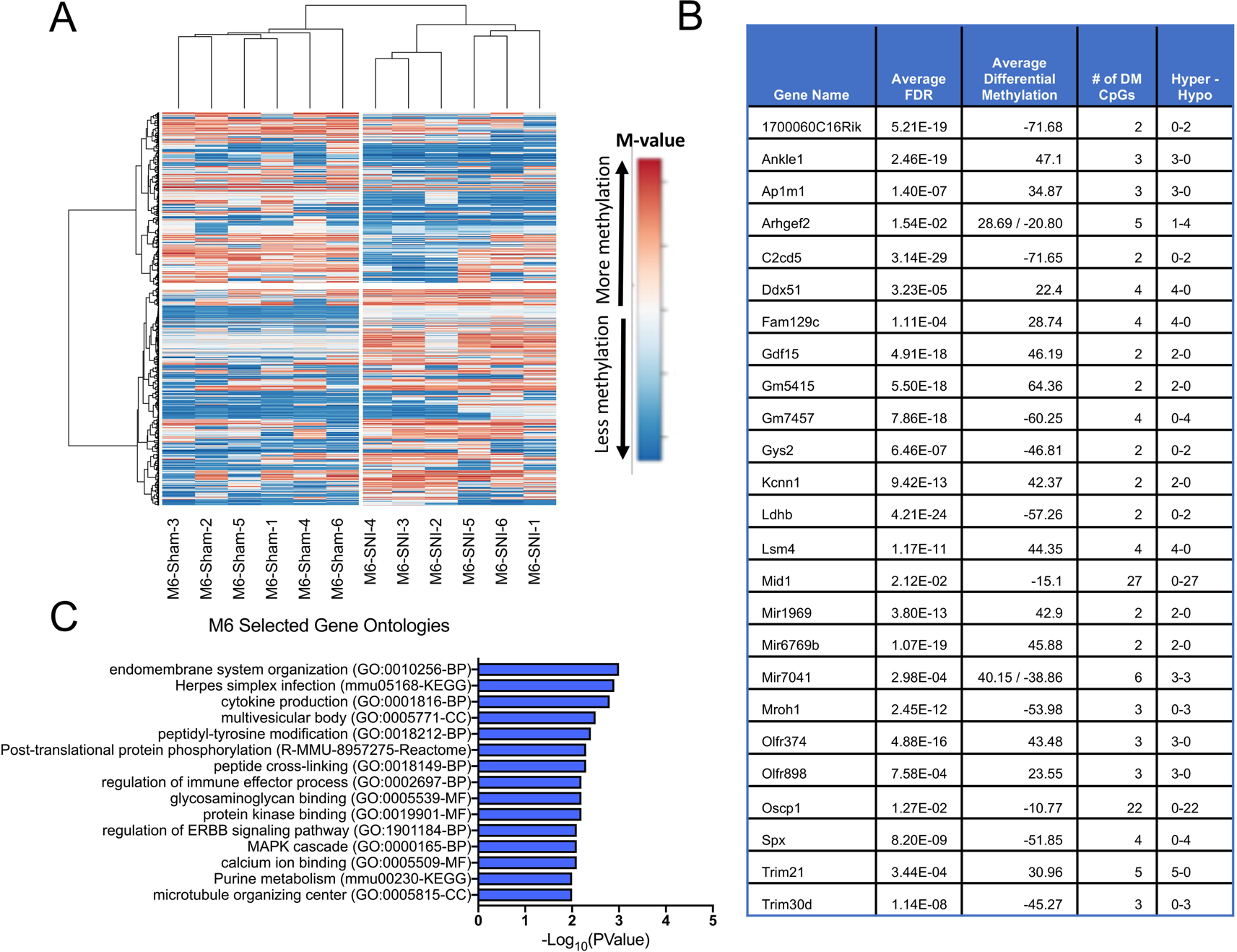
Differentially methylated genes and gene ontologies 6 months (6M) following spared nerve injury (SNI). **A)** Post-hoc supervised hierarchical clustering of DM CpGs at each time point displays common patterns of differential methylation between individual animals in the same treatment group. Values are displayed as M-Value: log_2_(# methylated CpGs / # unmethylated CpGs) Clustering method: Ward, distance metric: 1-Pearson correlation. **B)** Top 25 differentially methylated genes, **C)** Top 15 selected gene ontologies identified. *Enriched ontology: minimum gene count of 3, p < 0.05, and a minimum enrichment of 1.5. n = 6 mice/group*

At M6, the gene ontologies are related to immunological responses and systems, transcription factors, signaling cascades, and enzymatic processes. Specifically, top ontologies included: GO:0010256 – Endomembrane System Organization (p = 0.001, 26 genes), GO:0001816 – Cytokine Production (p = 1.58×10^-3^, 42 genes), GO:0018212 – Peptidyl-Tyrosine Modification (p = 3.98×10^-3^, 21 genes), GO:0002697 – Regulation of Immune Effector Process (p = 6.31×10^-3^, 25 genes), GO:0019901 – Protein Kinase Binding (p = 6.31×10^-3^, 38 genes), GO:0000165 – MAPK Cascade (p = 7.94×10^-3^, 40 genes), GO:0005509 – Calcium Ion Binding (p = 7.94×10^-3^, 33 genes), and GO:0005815 - Microtubule Organizing Center (p = 0.01, 35 genes)(**Figure 7C**).

### 3.4.4 One year post-injury

At 1Yr, 435 DM CpGs identified 359 unique genes undergoing differential methylation in their promoter region. The top 25 differentially methylated genes are involved in actin regulation (*Mprip, Parva, Txnrd1*), histone methylation (*Smyd1*), and the acid sensing channel *Asic3* (**Figure 8B**), In addition to the top 25, genes were identified that are involved with cell adhesion molecules (*Cldn13, Cldn15, Hapln3, Madcam1, Nectin2, Nfasc*) and extracellular matrix (*Col17a1, Lama3*).

**Figure 8.**
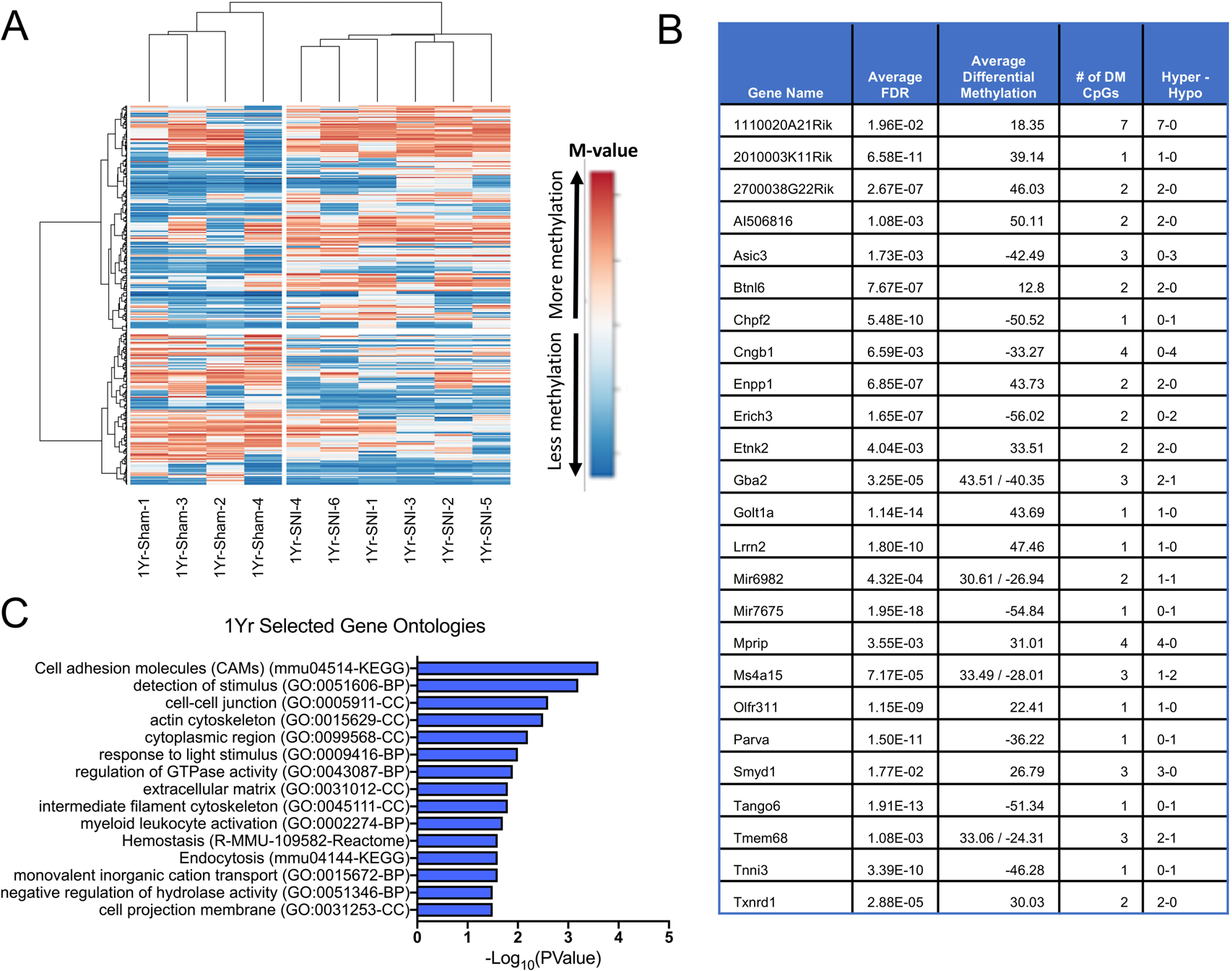
Differentially methylated genes and gene ontologies 1 year (1Yr) following spared nerve injury (SNI). **A)** Post-hoc supervised hierarchical clustering of DM CpGs at each time point displays common patterns of differential methylation between individual animals in the same treatment group. Values are displayed as M-Value: log_2_(# methylated CpGs / # unmethylated CpGs) Clustering method: Ward, distance metric: 1-Pearson correlation. **B)** Top 25 differentially methylated genes, **C)** Top 15 selected gene ontologies identified. *Enriched ontology: minimum gene count of 3, p < 0.05, and a minimum enrichment of 1.5. n = 4-6 mice/group*

At 1Yr, the top selected gene ontologies are related to cellular signaling, extracellular matrix formation, and cellular function regulation. Specifically, top ontologies include: mmu04514 – Cell Adhesion Molecules (CAMs) (p = 2.51×10^-4^, 9 genes), GO:0051606 – Detection of Stimulus (p = 6.31×10^-4^, 10 genes), GO:0005911 – Cell-Cell Junction (p = 2.51×10^-3^, 14 genes), GO:0005911 – Actin Cytoskeleton (p = 3.16×10^-3^, 14 genes), GO:0043087 – Regulation of GTPase Activity (p = 0.012, 9 genes), GO:0043087 – Regulation of GTPase Activity (p = 0.012, 9 genes), GO:0031012 – Extracellular Matrix, (p = 0.015, 12 genes), R-MMU-109582 – Hemostasis, (p = 0.025, 12 genes), and GO:0015672 – Monovalent Inorganic Cation Transport (p = 0.025, 11 genes) (**Figure 8C**).

## 3.5 Spared nerve injury results in function-specific responses across the time course

To identify trends in the functional responses to peripheral nerve injury across time, we overlapped all gene ontologies that were significantly enriched in differentially methylated genes at all time points. Overall, of the 775 enriched gene ontologies identified across all four time points, 658 gene ontologies were unique to one time point, 91 overlapped across two time points, 6 were found at three time points, and none were found to overlap across all four time points (**Figure 9A**). Displayed are the top 5 overlapping ontologies between time points, as determined by greatest averaged p-values and a minimum of 3 enriched genes at each time point in the ontology (**Figure 9B**). When identifying the overlapping gene ontologies at each time point, early time points D1 and W2 display ontologies more related to responding to external stimuli and damage, while later time points M6 and 1Yr display ontologies more related to adaptive immune response and T-cell mediated cytotoxicity. In all overlapping ontologies between time points, there is a strong contribution of actin and cytoskeleton related ontologies. These dysregulated pathways may contribute to functional and structural cortical remodeling previously reported in chronic pain conditions.

**Figure 9:**
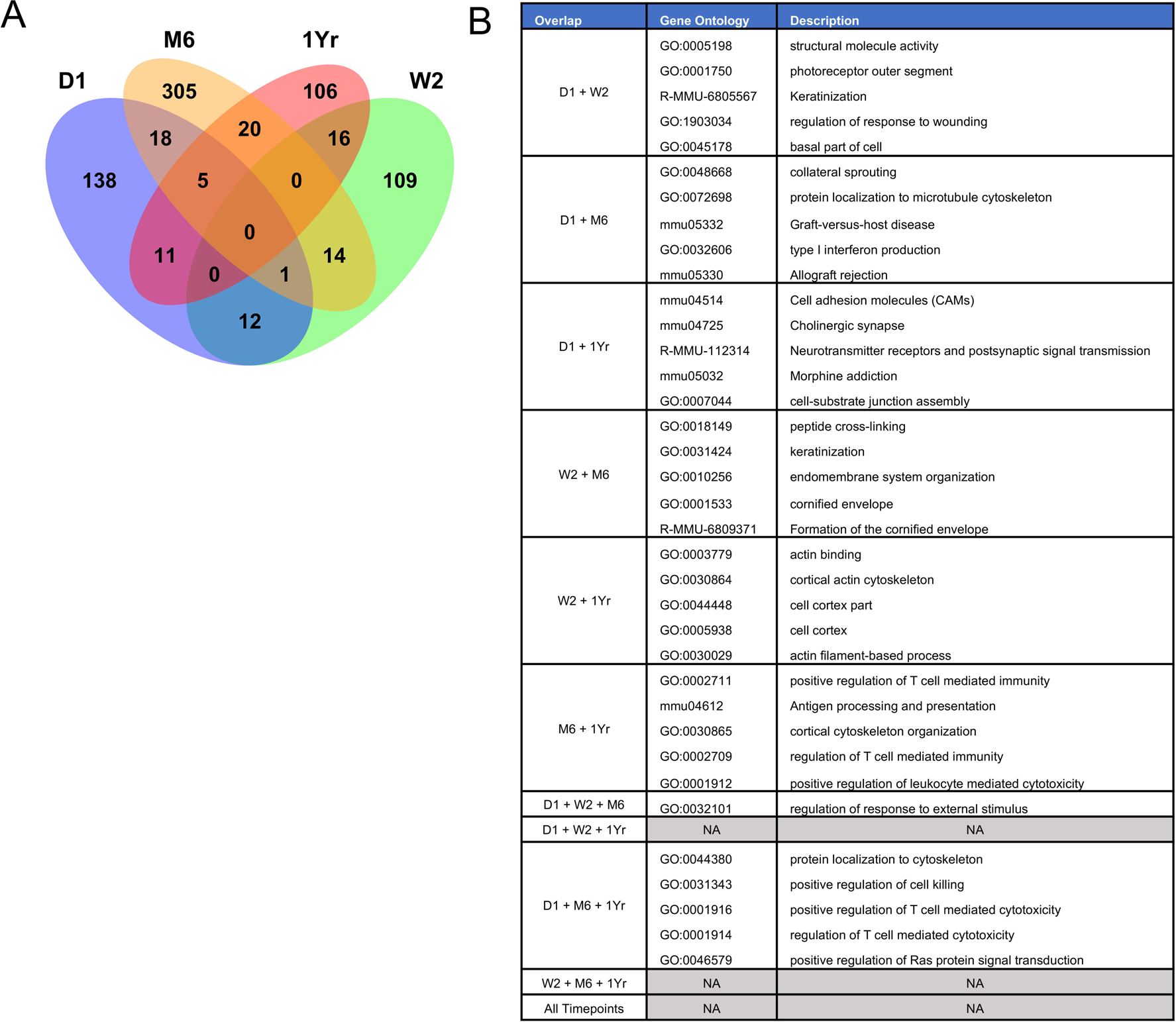
Overlap of dysregulated gene ontologies across time following spared nerve injury (SNI). **A)** Gene ontology analysis reveal that 658 of 775 identified enriched ontologies are unique to one time point, with only 91 ontologies overlapping at two or more time points. **B)** The top 5 gene ontologies per overlap as determined by greatest averaged p-value and a minimum of 3 enriched genes at each time point in the ontology

## 3.6 Differential methylation of pain-related genes show time point-specific patterns of differential methylation

To determine if any of the identified DM genes had been previously linked to pain, we cross-referenced with datasets of previously identified pain-related genes (**Figure 10**). Ultsch et al. [65] previously identified 535 genes associated with pain. After conversion to mouse gene homologs, we identified 31 differentially methylated genes from our experimental dataset. The Human Pain Genes Database (HPGD) has identified 361 genes in humans wherein SNPs have been found to affect the sensation of pain [47]. 289 HPGD genes had known mouse homologs, of which 15 differentially methylated genes were found in our dataset.

**Figure 10:**
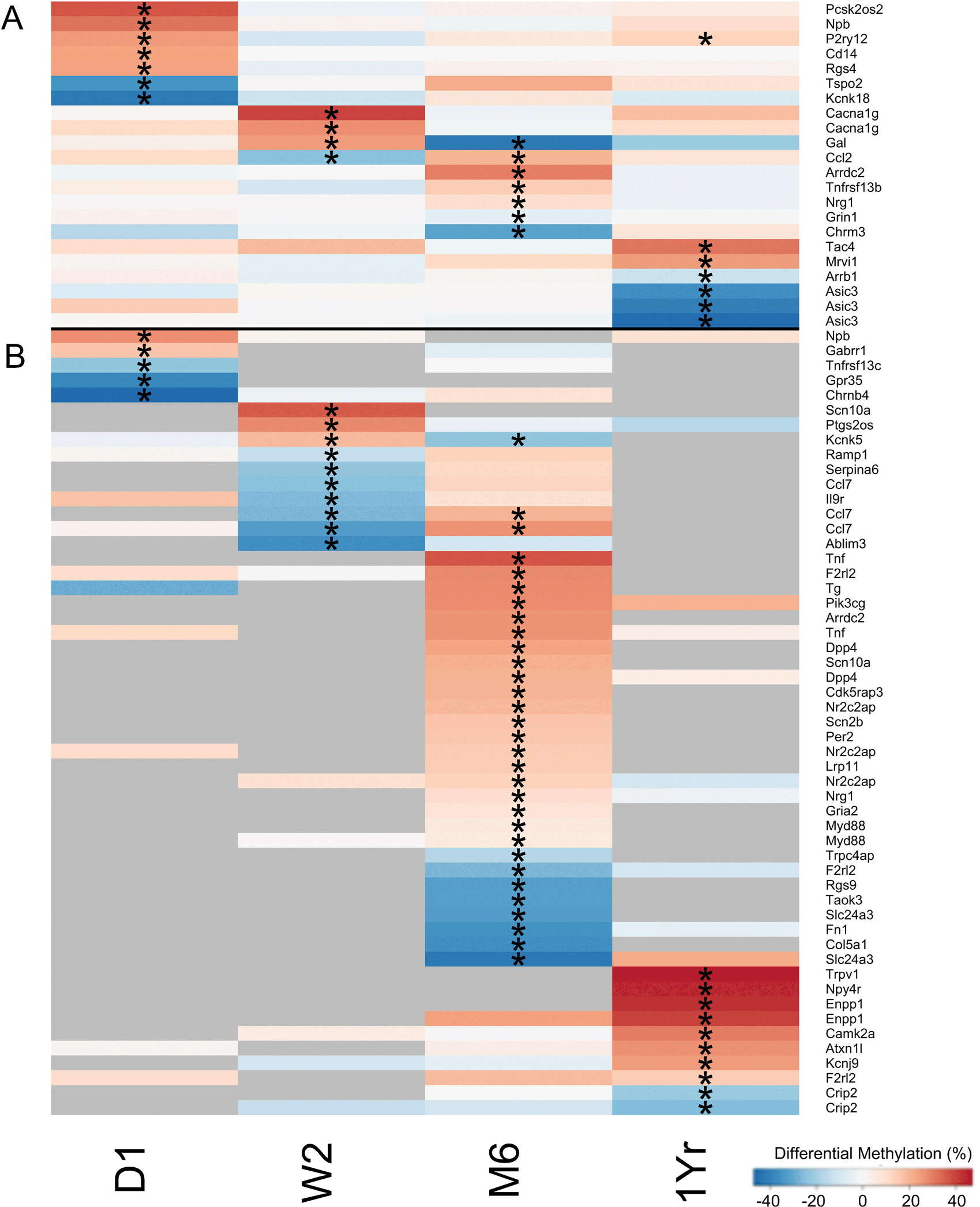
Identification and time course of differentially methylated CpGs associated with pain-related genes. Displayed are CpG positions that were annotated to a pain-related gene identified by Ultsch et al. [65] or the Human Pain Genes Database (HPGD) [47]. **A)** For CpGs that were captured at all time points, pain-related genes show mostly single time point differential methylation. **B)** For CpG positions that were not captured at all time points, there is a trend towards single time point differential methylation. Values are displayed as differential methylation. *\* = FDR adjusted p-value < 0.1. Grey = no data for that gene/time point*.

When tracking the differential methylation pattern of pain-related genes across the time course, most only show differential methylation at one time point. At D1, pain-related genes were involved in immune response (*Cd14, Tnfrsf13c*), and a number of ion channels (*Chrnb4, Gabrr1, Kcnk18*). At W2, pain-related genes were related to actin (*Ablim3*), immune regulation (*Ccl2, Ccl7, Il9r*), and ion channels (*Cacna1g, Kcnk5, Scn10a*) At M6, identified pain-related genes are involved in immune responses (*Ccl2, Ccl7, Dpp4, Myd88, Pik3cg, Tnf, Tnfrsf13b, Trpc4ap*), cell-cycle regulation (*Cdk5rap3, Taok3*), and ion channels and transport (*Chrm3, Kcnk5, Scn2b, Scn10a, Slc24a3*). At 1Yr, pain-related genes were involved in ion channels (*Asic3, Kcnj9,* Trpv1), peptide signaling (*Npy4r,* Tac4), and G-protein coupled receptor signaling (*Arrb1, P2ry12*).

For pain-related CpGs that were able to be captured at all time points, only three CpGs had differential methylation at more than one time point (*P2ry12, Gal,* and *Ccl2*) (**Figure 10A)**. Interestingly, galanin (*Gal)* and chemokine ligand 2 (*Ccl2)*, both show reversals of their methylation state from W2 to M6. The methylation reversal of these two pain genes may indicate that the recruitment of pain-related processes may change across the time course, and genes have a temporal impact upon chronic pain development. While limited by a lack of data at all time points, a similar time point specific pattern of significant differential methylation was observed for CpGs that were not captured at all time points (**Figure 10B).** Therefore while a gene may be pain-related, its contribution to a pain phenotype may be restricted or limited in time.

We also cross-referenced our experimental genes with two gene sets identified in our previous studies. Massart et al. 2016 examined rat PFC DNA methylation at 9 months post-SNI and identified 3981 genes that undergo differential methylation [43]. Alvarado et al. 2013 examined mouse PFC mRNA expression using RNAseq at 6-8 months post-SNI and identified 641 differentially expressed genes [1]. After conversion to mouse homologs, 76 genes overlapped between M6 genes and Massart et al., 39 genes displayed consistent differential methylation profiles between the two species (Supplemental Table 5). 27 genes overlapped between M6 genes and Alvarado et al., with 12 differentially methylated genes consistent with the predicted change in mRNA expression on (Supplemental Table 6).

The NMDA receptor subunit *Grin1* was identified in all three datasets, showing both consistent methylation across species and the predicted impact on mRNA expression. Gene regulatory genes *Eif3g* and *Dpy30*, and neurite outgrowth inhibitor *Rtn4* were among the genes consistent between M6 genes and Massart et al. Among the gene consistent between M6 and Alvarado et al. were the cell-cell junction desmosome gene *Dsg1c*, the actin regulatory gene *Cobl*, and the calcium/calmodulin dependent protein kinase *Camk1d*.

## 4. Discussion

This study evaluated DNA methylation as a mechanism for the long term embedding of an injury-related pain state in the PFC and the subsequent evolution of chronic pain. Previous studies have demonstrated a DNA methylation response to chronic pain in the PFC of rodents [43; 62], however a number of questions still remain. Is DNA methylation a consequence of chronic pain or do these changes precede its emergence? How rapidly do DNA methylation changes occur, and are they persistent throughout the time course of chronic pain? How targeted are these changes? Are they limited to pain genes, or do they affect entire domains and gene families? What can these changes tell us about the mechanisms of how chronic pain develops and embeds itself? Finally, and most importantly, which of these processes should be targeted for intervention?

To answer these questions we tracked PFC DNA methylation profiles for one year post-SNI. We provide first evidence that the PFC methylome is reprogrammed immediately after injury at D1, supporting the hypothesis that DNA methylation changes precede the emergence of chronic pain. Second, we show that SNI triggers a cascade of dynamic and evolving DNA methylation changes that continue up to one year post-injury, suggesting that DNA methylation changes accompany the onset and establishment of chronic pain phenotypes. Third, we identified hundreds of genes and numerous gene ontologies that experience differential methylation. While some of these may be relevant at only one time point, others are differentially methylated throughout the time course, potentially serving as the genomic memory of the initial injury and necessary for chronic pain development.

### 4.1 CpG methylation dynamics

We first analyzed differential CpG methylation between Sham and SNI at each time point independently. At one day post-injury, we observed the differential methylation of hundreds of specific CpG positions in the PFC. The D1 time point had the second largest proportion of differentially methylated CpGs identified and the largest average differential methylation, illustrating that the response of the PFC methylome to an injury is rapid and substantial. Each time point evaluated had a large population of DM CpGs that display a substantial significant difference (FDR adjusted p-value < 1E-7) and large changes in differential methylation (> 40%), indicating strong methylation changes take place throughout the chronic pain time course.

The presence of both hyper-methylated and hypo-methylated CpG positions at each time point implies targeted regulation rather than a genome-wide increase or decrease in methylation. There is also a slight trend towards overall promoter hypermethylation, which may indicate a reduction in overall rates of transcription or a greater number of silenced genes. We previously observed that global DNA methylation decreases after SNI [62], however this difference can be explained by the non-specific demethylation of non-promoter regions in the genome.

### 4.2 Differentially methylated genes, pathways, and functions

We identified 1633 total unique genes that experience differential promoter methylation, of which 31 and 15 genes were members of the pain-related gene lists by Ultsch et al. or Human Pain Genes Database respectively. Thus, chronic pain effects not just a few candidate genes but also functionally related groups of genes. Each time point had at least 300 differentially methylated genes identified, and over 150 enriched gene ontologies. At the early time points of D1 and W2, we observed a greater number of ontologies involved in the response to external stimuli, while at the later time points of M6 and 1Yr, inflammatory processes and T-cell adaptive immune responses are implicated. Interestingly, across all four time points numerous ontologies involved in actin dynamics or cytoskeletal regulation were identified. Actin reorganization has been previously identified as a key step in inflammation related persistent pain in glial cells in the spinal cord [25], and sensory neurons in the periphery [16]. The reorganization of nociceptive circuits and dendritic pruning in response to peripheral neuropathic pain is well established [48; 63; 64], and may be a form of maladaptive synaptic plasticity. The consistent differential methylation of actin and cytoskeleton related genes throughout the time course for up to one year post-injury suggests an ongoing cortical plasticity and remodeling in response to peripheral injury, and that chronic pain induces progressively greater deviations from normal cortical functioning.

Interestingly, the NMDA receptor subunit Grin1 was differentially methylated in both mouse and rat PFC 6 and 9 months post-SNI, respectively and Grin1 mRNA expression was dysregulated in mouse cortex during the same time frame [1; 43]. Grin1 has a well-established role in pain-related neuroplasticity and these converging lines of evidence suggest that its dysregulation in chronic pain may be epigenetically-mediated. The presence of consistently differentially regulated genes across different studies using different animal models or focusing on different regulatory mechanisms may help to identify crucial pain-related genes.

### 4.3 Time point specific methylation patterns across the chronic pain time course

DM CpGs follow different trajectories of differential methylation across the progression from acute to chronic pain. These differential methylation profiles are a time stamp for the contributions of each gene during chronic pain development and shed light on the post-injury response. Genes with DM CpGs that maintain a methylated state throughout the time course, such as *Wipf2, Rrp8, Xdh, or Sprr2d* may encode a persistent genomic memory of the initial injury, triggering and maintaining a chronic pain state.

However, equally important to consider is the time point specific differential methylation of genes. Nearly all identified pain-related genes were differentially methylated at a single time point, implying a time-dependent recruitment or dysregulation of normal functioning. *Grin1, Nrg1, and Asic3* had confirmed single time point differential methylation, and while *Gabrr1, Scn10a, Ramp1, Myd88, Trpv1, Tnf* were unconfirmed, all experienced significant differential methylation and have well documented impacts on pain sensation across domains of ion channel activity and trafficking, inflammatory responses, and neurotransmitter regulation [15; 17; 26-28; 31; 39-42; 44; 45; 61; 66; 73].

The time point specific gene differential methylation and ontology enrichment underscore the importance of understanding temporal dynamics and effects upon normal functioning during the transition to chronic pain. For example, we identified 32 differentially methylated genes involved T-Cell activation, however these genes are spread throughout the time course and can be differentially methylated at one time point or multiple (**D1**: *Adam17, Btnl6, Cxcr4, Dusp3, Icos, Itch, Rhoh, Rpl22, Tnfrsf13c, Txk*; **W2**: *Btla, Btnl6, Ccl2, Cxcr4, Rhoh*; **M6**: *Ankle1, Ccl2, Ccnd3, Cd6, Dpp4, Dusp3, Il12b, Il12rb1, Jak3, Pla2g2e, Pla2g2f, Prex1, Ptpn22, Rpl22, Spta1, Tcf7, Tcirg1, Traf6, Zap70;* **1Yr**: *Btnl6, Cd6, Gimap1, Il1f8, Il27, Rpl22, Tcf3*).

While some of the observed DNA methylation changes may be collateral to chronic pain responses, many are likely to be required to initiate the chronic pain phenotype. The involvement of multiple genes poses a significant challenge for reversing chronic pain. Broader interventions such as targeting nodal regulatory genes such as DNMTs or other epigenetic machinery might be an alternative strategy for reversing this complex and multigenic epigenetic rearrangement.

### 4.4 Limitations

First, only male mice were studied. Given known sex differences in pain processing [60], our results lack the female methylation response to chronic pain development, especially important given identified sex differences in aging-related cognition and genetics [18; 24]. Second, our experiment focused on promoter region methylation, potentially missing DNA methylation regulation in non-promoter regions or by other processes. DNA methylation in the first intron of coding genes or gene body can affect gene expression [3; 71]. Moreover, DNA methylation does not have a linear relationship to gene expression, with other regulatory processes such as RNA transcription or protein translation involved, thus requiring a future analysis of gene expression. Third, our tissue samples are derived from whole brain tissue homogenates, and therefore mixtures of different brain cell types, each with their own cell type specific profiles of DNA methylation. This presents the possibility for differential methylation signal to be diluted by conflicting methylomic data. However, given the lack of significant changes in major cell type proportions (Figure 2), this impact is likely minimal. Fourth, the alpha threshold was set at an FDR of 0.1 in this study. It is acknowledged that the selection of an FDR threshold of 0.1 carries a greater risk of type I error (false positive), it is preferable for the exploratory purposes of this study where type II error (false negative) is a greater concern. We recognize these limitations and present these results as a first step towards understanding the epigenetic reprogramming associated with the development and maintenance of chronic pain.

## 5. Conclusions

Our findings provide evidence for rapid epigenetic reprogramming in the PFC methylome in response to peripheral nerve injury. Reprograming persists as pain becomes chronic, thereby suggesting an important role for DNA methylation in the initiation, progression and maintenance of chronic pain. While some changes in DNA methylation were persistent, dynamic reprogramming was also observed. These patterns may reflect underlying differences between acute, sub-chronic and chronic pain, suggesting that therapeutic interventions may have efficacy dependent on pain duration. Pathway analysis revealed enrichment in genes related to stimulus response at early time points, immune function at later time points and actin and cytoskeletal regulation throughout the time course. These findings highlight the importance of understanding the time course of acute to chronic pain and suggesting broader pain interventions than single gene or protein targets are required.

## Acknowledgements

The authors would like to thank the Louise and Alan Edwards Foundation, the Canadian Institutes of Health Research, and the Fonds de la Recherche en Sante du Quebec for their financial support. The authors would also like to thank the McGill Animal Resources Centre, Compute Canada, the McGill University Genome Quebec Innovation Centre, and Virginie Calderon, Odile Neyret, Francois Lefebvre, and Xiaojian Shao for valuable advice and technical support. The authors declare no conflicts of interest.

## Figure Legends

**Supplemental Figure 1.**
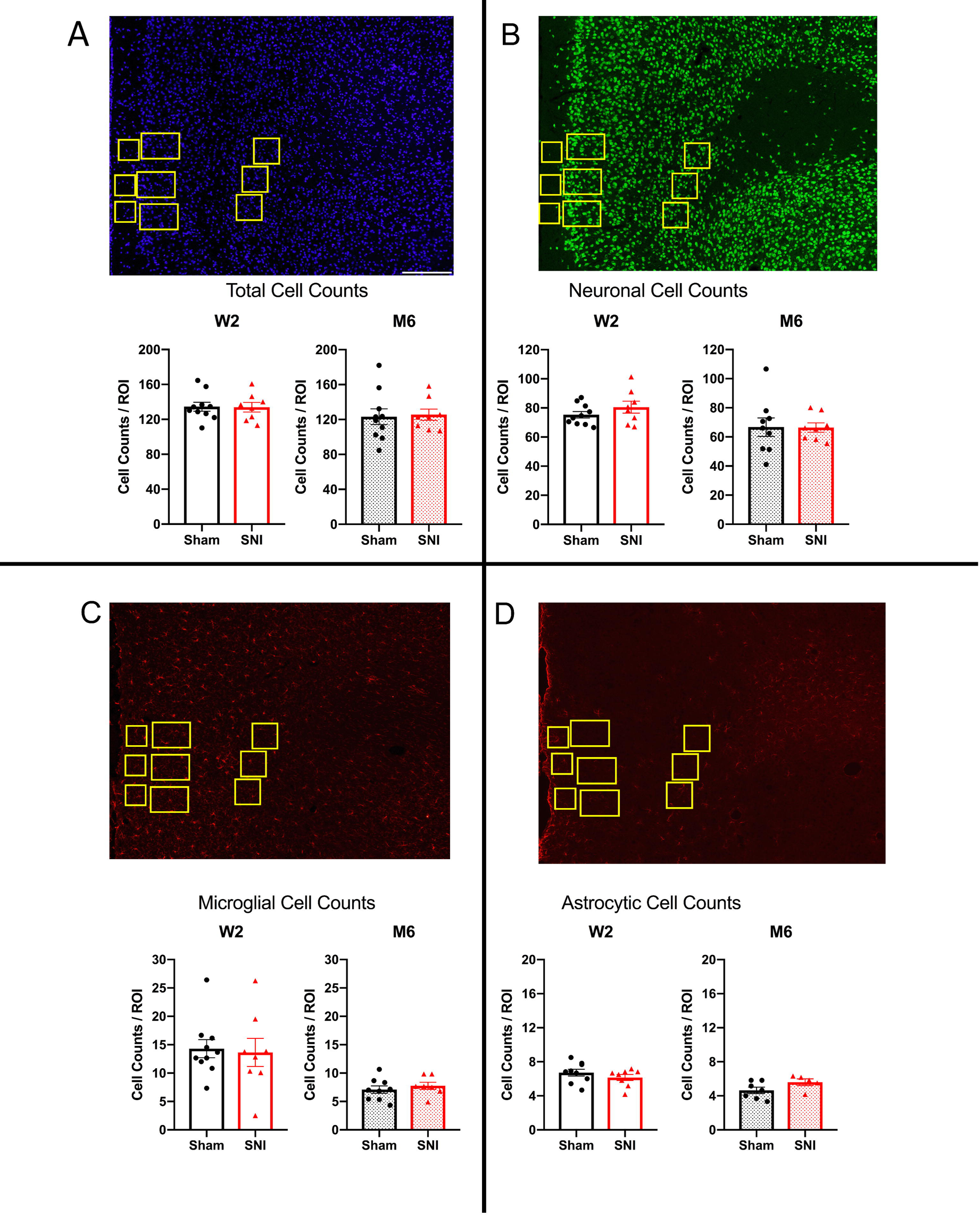
Immunohistochemistry reveals no changes in neuronal, microglial or astrocytic cell proportions two-weeks and six-months in infralimbic cortex following spared nerve injury (SNI). A) Total (DAPI, Blue), B) Neuronal (NeuN-immunoreactivity (-ir), Green), C) microglial (Iba1-ir, Red), and D) astrocytic (GFAP-ir, Red) cell counts in Sham and SNI cortex at W2 or M6 time points. Displayed are average cell counts from all cortical layers from prelimbic area of the infralimbic cortex. Quantified ROI area is equivalent to 33040.9 µm^2^. Bars represent mean ± SEM cell counts; *Two-tailed Welch’s t-test. No significant differences observed. n = 8-10/group*.

**Supplemental Table 1 – Probe Bed File.** Genomic coordinates for the bisulfite capture-sequencing probes

**Supplemental Table 2 – Blacklist Bed File.** Genomic coordinates for regions known to have anomalous, unstructured, and high signal/read counts in next gen sequencing

**Supplemental Table 3 – All differentially methylated CpGs and annotated genes identified**

**Supplemental Table 4 All enriched gene ontologies identified**

**Supplemental Table 5.**
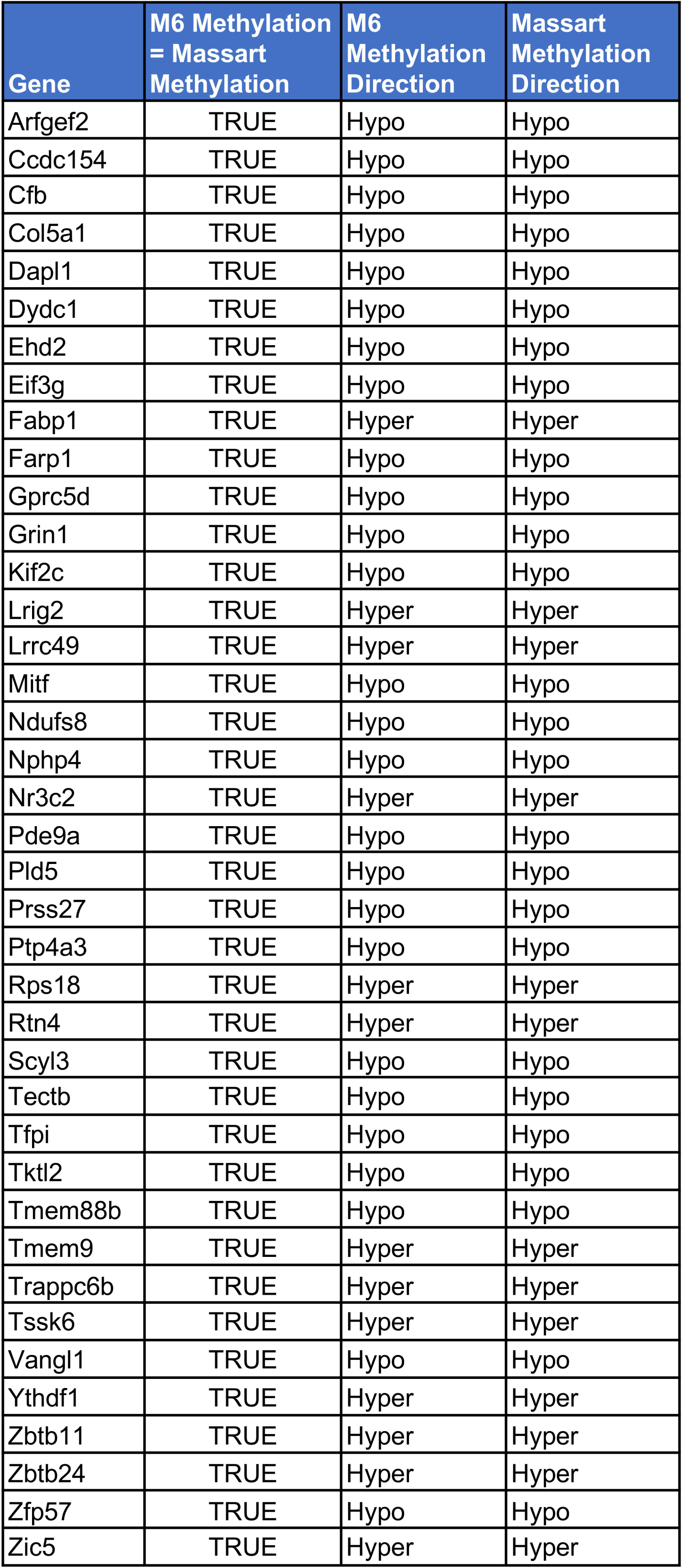
Genes with consistent differential methylation profiles between M6 and those identified previously in Massart et al. 2016

**Supplemental Table 6.** Genes consistent with the predicted change in mRNA expression between M6 and those identified previously in Alvarado et al. 2013

| Gene | M6 Methylation<br>= Alvarado<br>RNASeq | Methylation<br>Direction | RNA<br>Transcription |
| --- | --- | --- | --- |
| Camk1d | TRUE | Hypo | Increased |
| Cobl | TRUE | Hypo | Increased |
| Cst3 | TRUE | Hyper | Decreased |
| Ddhd1 | TRUE | Hypo | Increased |
| Dsg1c | TRUE | Hyper | Decreased |
| Grin1 | TRUE | Hypo | Increased |
| Hnrnpab | TRUE | Hypo | Increased |
| Kcnip2 | TRUE | Hypo | Increased |
| Rps21 | TRUE | Hyper | Decreased |
| Serf2 | TRUE | Hyper | Decreased |
| Snhg11 | TRUE | Hyper | Decreased |
| Tmem201 | TRUE | Hypo | Increased |

